# *Escherichia coli* populations adapt to complex, unpredictable fluctuations by minimizing trade-offs across environments

**DOI:** 10.1101/070318

**Authors:** Shraddha Karve, Devika Bhave, Dhanashri Nevgi, Sutirth Dey

## Abstract

In nature, organisms are simultaneously exposed to multiple stresses (i.e. complex environments) that often fluctuate unpredictably. While both these factors have been studied in isolation, the interaction of the two remains poorly explored. To address this issue, we selected laboratory populations of *Escherichia coli* under complex (i.e. stressful combinations of pH, H_2_O_2_ and NaCl) unpredictably fluctuating environments for ~900 generations. We compared the growth rates and the corresponding trade-off patterns of these populations to those that were selected under constant values of the component stresses (i.e. pH, H_2_O_2_ and NaCl) for the same duration. The fluctuation-selected populations had greater mean growth rate and lower variation for growth rate over all the selection environments experienced. However, while the populations selected under constant stresses experienced trade-offs in the environments other than those in which they were selected, the fluctuation-selected populations could by-pass the across-environment trade-offs almost entirely. Interestingly, trade-offs were found between growth rates and carrying capacities. The results suggest that complexity and fluctuations can strongly affect the underlying trade-off structure in evolving populations.

## INTRODUCTION

Experimental evolution using microbes has led to a large number of insights in evolutionary biology (reviewed in Kassen, 2014). Much of those insights pertain to microbial populations evolving under a single selection pressure which is either static or changing directionally (reviewed in Scheiner, 2002; Lenski, 2004). However, in nature, multiple selection pressures typically not only operate simultaneously, but also undergo predictable or unpredictable spatiotemporal fluctuations. Theoretical studies over the last few decades have indicated that the evolutionary outcomes of selection under such conditions can be very different from exposure to directional selection under a single environment (reviewed in Levins, 1968; Whitlock, 1996).

Theoretical studies predict that the response to fluctuating environments depend on the rate at which the environments change. Thus, in asexual populations, phenotypic plasticity evolves when the environments change very rapidly, i.e. within few generations (Ancel, 1999) but evolution by selective sweeps of large-effect mutations is favoured when the fluctuations occur after hundreds of generations (Cohan, 2005). Another way by which populations can evolve under temporal fluctuations is by reducing the variance of fitness over time (or across changing environments), through bet-hedging (Holman, 2015 and references therein). Although there is some controversy in terms of its usage in the experimental evolution literature (de Jong *et al.*, 2011), empirical evidences suggest that bet-hedging does evolve in microbes in response to environmental fluctuations (Veening *et al.*, 2008; Ratcliff & Denison, 2010; Graham *et al.*, 2014). Interestingly, although there are quite a few insights on evolutionary responses to very fast and very slow rates of environmental changes, the effects of intermediate rates of changes (i.e. on the time scale of tens of generations) have proven difficult to predict and are thought to critically depend on the underlying genetic architecture (Chevin *et al.*, 2010).

Another crucial aspect of temporal fluctuations that seems to play a major role in determining the evolutionary dynamics of the populations is predictability. Environmental fluctuations can be predictable like seasonal temperature variations, or unpredictable like the chemical parameters of a sewer carrying domestic sewage. In case of asexual organisms, predictable within-generation fluctuations are expected to select for switching phenotypes where the rate of switching evolves to match the rate of fluctuations (Acar *et al.*, 2008). Unpredictable fluctuations on the other hand, are likely to favour bet-hedging with random phenotypic switch (Beaumont *et al.*, 2009). Similarly, there have been a number of empirical studies on the fitness outcomes of different kinds of environmental fluctuations (Leroi *et al.*, 1994; Reboud & Bell, 1997; Hallsson & Björklund, 2012; Alto *et al.*, 2013; Ketola *et al.*, 2013; Condon *et al.*, 2014; Karve *et al.*, 2015; Karve *et al.*, 2016). For example, microbial populations facing predictable environmental fluctuations show improved fitness over the entire range of fluctuating environments (Leroi *et al.*, 1994; Hughes *et al.*, 2007; Coffey & Vignuzzi, 2011). On the other hand, unpredictable fluctuations sometimes improve fitness of the microbes in all environments (Turner & Elena, 2000; Ketola *et al.*, 2013) or improve in some of the environments without any change in others (Hughes *et al.*, 2007).

Another aspect that characterizes natural environments is complexity, i.e. the simultaneous presence of multiple selection pressures. Unfortunately, this aspect of microbial evolution remains relatively unexplored with almost no theoretical studies and few empirical investigations. Complex environments have been shown to result in fitness improvements, albeit to different degrees, for all the resources experienced during selection (Barrett *et al.*, 2005; Cooper & Lenski, 2010). Unlike mean fitness though, variation for fitness shows opposing trends for fluctuating and complex environments. Populations selected under fluctuating values of a single environment (say different temperatures) often show reduced variation for fitness over the entire range of values experienced during selection. (Kassen, 2014 Table 4.2) However when populations are selected under environments containing multiple parameters (say different carbon sources), they tend to have highly variable fitness when assayed under various media containing only one of those parameters (Barrett *et al.*, 2005).

Thus, although there is some understanding of how microbes adapt to environmental fluctuations and complexity, the evolutionary outcomes of simultaneously facing environmental fluctuations and complexity remain relatively unexplored in terms of mean fitness (although see Karve *et al.*, 2015) as well as variation for fitness. This lack of understanding also extends to the underlying constraints experienced by evolving populations, particularly in the context of the patterns of trade-offs.

In life-history evolution, a trade-off represents a scenario wherein a beneficial evolutionary change is accompanied by a detrimental one (Stearns, 1989). This can happen in at least two different ways. Trade-off across traits happens when, for a given environment, an increase in fitness *w.r.t.* some trait(s) coincides with a decrease in fitness *w.r.t.* some other trait(s) (Roff & Fairbairn, 2007). Alternatively, trade-off across environments take place when, for a given trait, an increase in fitness in some environment(s) is accompanied by a reduction in fitness in some other environment(s) (Anderson *et al.*, 2014). Across environments trade-offs are typically what is studied in microbial systems and are also of interest in the context of this study. This is because, in such systems, it is often difficult to separately estimate the various components of fitness in a given environment. This is also the reason for which most fitness measurements in microbial populations involve composite traits like growth rate or competitive ability. Interestingly, in microbial experimental evolution studies, trade-offs are mostly observed in novel environments, i.e. environments not experienced during selection (Bennett *et al.*, 1992; Reboud & Bell, 1997; Hughes *et al.*, 2007; Jasmin & Kassen, 2007) and rarely across selection environments. However, little is known about the trade-off structure in microbial populations when they are exposed to multiple selection environments changing unpredictably.

Here we study the evolutionary implications of the interaction of complexity and temporal fluctuations in environments. Replicate laboratory populations of *Escherichia coli* were exposed to complex (i.e. stressful combinations of pH, H_2_O_2_ and NaCl), unpredictably fluctuating environments. Parallely, we also selected replicate bacterial populations under constant exposure to each of the selection environments. After ~900 generations of selection, we found that populations facing complex, unpredictable environments increased their mean fitness (measured as population growth rates) and minimized the variation for fitness when estimated over all the selection environments. Moreover, selection under complex, fluctuating environments did not lead to trade-offs across different selection environments, although some trade-offs were observed between growth rate and carrying capacity in some of the environments. Our results show that populations facing unpredictably fluctuating complex environments can show higher mean fitness and significantly lower fitness variation across environments, without undergoing any trade-offs across selection environments.

## METHODS

### Selection experiment

We used Kanamycin resistant *Escherichia coli* strain K12 (see Supplementary online material S1 for details) for this study. A single colony grown on Nutrient agar with Kanamycin (see SOM for composition) was inoculated in 2 ml of Nutrient broth with Kanamycin (NB^Kan^) (see Supplementary online material S2 for composition) and allowed to grow for 24 h at 37°C, 150 rpm in 24 welled plate. 4 μl of this suspension was used to initiate each of 120 replicate populations.

120 replicate populations were equally divided into five treatment regimes and one control regime, such that there were 20 replicate populations per regime. Control populations were subcultured in NB^Kan^ for the entire duration of the selection. Four out of five selection regimes were constant environments with salt or pH9 or pH4.5 or hydrogen peroxide (H_2_O_2_) in NB^Kan^. The remaining selection environment was complex and stochastically fluctuating (henceforth termed as F) (see Supplementary online material S3 for details of all selection regimes). Briefly, we created 64 arbitrary combinations of NaCl, pH and H2O2 values such that in any combination, the magnitude of one component was equal to that found in NB (i.e. pH = 7.0 or [NaCl] = 0.5g% or [H2O2] = 0) and that of the other two were individually expected to negatively affect growth. Thus, for example, combination # 46 denotes a NB containing 3.5 g% of NaCl + 7pH + 3.4 mM H_2_O_2_ whereas combination # 10 stands for 4.5 g% of NaCl + 4.5 pH + no H_2_O_2_ (Supplementary material S3). Each combination was assigned a tag between 0-63 and a uniform-distribution random number generator was used to obtain a sequence of numbers (with replacement) in that range. Each replicate F population experienced 100 different combinations of the components according to this sequence. This ensured that the environments faced by these populations were not only unpredictable, but also complex, i.e. involved three different environmental variables. The details of this fluctuating selection regime has been mentioned elsewhere (Karve *et al.*, 2015).

24 welled plates with 2 ml of appropriate growth medium and 4 μl of inoculum volume for each well were used throughout the selection and assay experiments. The growth conditions were maintained at 37°C, 150 rpm. All the populations were sub-cultured every 24 h. Extinctions were identified visually as a lack of turbidity and revived using 20 μl of the previous day’s culture stored at 4°C. The selection lasted for 100 days i.e. ~ 900 generations, computed using the standard expression for calculation of generation time (Bennett & Lenski, 1997). On the100^th^ day, the populations were stored as glycerol stocks at -80°C for future assays.

### Fitness assay in selection environments

After 100 days of selection, all the populations were assayed for fitness in every selection environment, except the fluctuating environment. Selection environments comprised of NB^Kan^ with salt or pH9 or pH4.5 or hydrogen peroxide or control (see Supplementary online material S3 for details). For comparisons with the ancestor, we revived the ancestral population of Kanamycin resistant *Escherichia coli* strain K12 in NB^Kan^ for the duration of 18 h. 20 replicate wells were inoculated with this revived culture for every selection environment. This resulted in the same number of replicates of ancestral culture for every assay environment as that of the selected populations.

We used two population-level measures of fitness, namely maximum growth rate and carrying capacity, both of which have individual level correlates. Population level growth rate correlates with the speed of growth and division of individual cells whereas carrying capacity correlates with the minimum amount of nutrients (or environmental conditions) needed for division or tolerance to the detrimental effects of density. Both these traits can potentially trade-off at the level of individuals and such trade-off can be reflected in the correlated population level estimates. Following previous studies, maximum growth rate during 24 h of growth was used as a fitness measure (Ketola *et al.*, 2013; Karve *et al.*, 2015). For the growth rate measurement, 4 μl of relevant glycerol stocks were revived in 2 ml of in NB^Kan^. After 18 h of growth, these revived cultures were inoculated in the appropriate assay environment. OD_600_ was measured every 2 h on a plate reader (Synergy HT BioTek, Winooski, VT, USA) for the duration of 24 h. We used a QBASIC (v 4.5) script to determine the maximum growth rate of the bacterial populations. The program fits a straight line on overlapping moving windows of three points on the time series of OD_600_ values. The maximum slope obtained by this method was taken as the maximum growth rate for that population.

We used the maximum density reached during the same 24 h growth trajectories as an estimate of the carrying capacity. Both fitness estimates, i.e. maximum growth rate and maximum density reached, were analyzed as mentioned below.

### Replicates and statistical analysis

For every population (6 selection regimes × 20 replicates + 20 replicates of the ancestral population) growth assays were performed twice in five environments (pH4.5, pH9, H_2_O_2_, salt, NB). This resulted in a total of 1400 growth measurements (140 populations × 5 assay environments × 2 measurements).

#### Overall mean fitness

1200 fitness estimates (6 selection regimes × 5 assay environments × 20 replicates × 2 measurements, excluding the ancestor) were analysed using a mixed-model with two fixed factors (selection with six levels and assay environment with five levels) and a random factor (replicate with twenty levels) nested in selection. This design allowed us to study how, for a given selection environment comprised of 20 replicate populations, the mean fitness differ across the assay environments. To determine whether F populations differed significantly from the other selection lines, we performed Dunnett’s post hoc test (Zar, 1999).

#### Variation for fitness

We estimated coefficients of variation (CV) as a measure of variation in fitness across all the assay environments. Two fitness estimates in a given assay environment were averaged and CV was calculated for every replicate population over this average fitness in five assay environments. Every selection regime thus yielded 20 CV estimates (one for each replicate population) which reflected the variation in fitness across five environments for the populations selected in that regime. These were then analyzed using a one way ANOVA with selection (six levels) as a fixed factor. To determine whether F populations differ significantly from other selection lines, we performed Dunnett’s post hoc test (Zar, 1999).

#### Trade off

The fitness of the ancestor was assayed in every environment used during the selection (i.e. NB, salt, pH 4.5, pH 9 and H_2_O_2_) and the mean fitness for a given environment was computed as the average of 40 values (20 replicates × 2 independent measurements). This was then subtracted from the mean fitness of every population assayed in that environment. Thus, for example, the mean fitness of the ancestor in salt was subtracted from the fitness values of all the selected populations (6 selection regimes × 20 replicates) assayed in salt. This resulted in 20 difference measurements for every selection regime in every environment. When the mean of these 20 values were significantly greater than zero, we interpreted that the corresponding selection regime has gained fitness in that assay environment and vice versa. To ascertain this, we performed 30 different ANOVAs (6 selection lines × 5 assay environments) and controlled for family-wise error rates through sequential Holm-Šidàk correction of the p-values (Abdi, 2010).

To estimate the biological significance of the difference in fitness of F populations compared to other selected populations, we computed Cohen’s *d* statistics (Cohen, 1988) as a measure of effect size. It was interpreted as small, medium and large for 0.2 < d < 0.5, 0.5 < d < 0.8 and d > 0.8, respectively.

All the ANOVAs were performed on STATISTICA v7.0 (Statsoft Inc.). Cohen’s *d* statistics were estimated using the freeware Effect size generator v2.3.0 (Devilly, 2004).

## RESULTS

### Fluctuating environments select for higher overall mean fitness and lower variation for fitness across environments

When analyzed over all the selection regimes, effect of selection was highly significant (*F*_6,700_ = 12.07, *p* < 0.0001). The effect of assay environment was also highly significant (*F*_4,700_ = 247.04, *p* < 0.001) and there was a significant interaction of selection with assay environment (*F*_24,600_ = 31.73, *p* < 0.001). The growth rates of the selected populations vis-a-vis the ancestor were compared using Dunnett’s test. F populations showed the highest overall mean fitness which was significantly greater than the mean fitness of ancestor with medium effect size (Table 1, Fig 1A). None of the other selected populations differed significantly from the ancestor.

**Table 1.** Summary of the main effect of selection in the ANOVAs for individual selection regimes. Dunnett’s post hoc test was conducted with ancestor as a control group. Dunnett’s test *p* values and effect size are thus in comparison with ancestor.

| **Assay** | **Selection regime** | **Mean** | **Dunnett *p* values** | **Effect Size±95% CI** | **Interpretation** |
| --- | --- | --- | --- | --- | --- |
| Overall mean fitness | Ancestor | 0.125 | - | - | - |
|  | pH4.5 | 0.129 | 0.826 | 0.07±0.19 | Small |
|  | pH9 | 0.114 | 0.016 | 0.2±0.19 | Small |
|  | NB | 0.125 | 1 | 0.002±0.19 | Small |
|  | H<sub>2</sub>O<sub>2</sub> | 0.133 | 0.185 | 0.14±0.19 | Small |
|  | Salt | 0.133 | 0.278 | 0.17±0.19 | Small |
|  | F | 0.146 | 0.00003 | 0.5±0.19 | Medium |
| Coefficient of variation for mean fitness | Ancestor | 0.284 | - | - | - |
|  | pH4.5 | 0.367 | 0.007 | 1.13±0.67 | Large |
|  | pH9 | 0.491 | 7.0E-06 | 1.6±0.82 | Large |
|  | NB | 0.357 | 0.02 | 0.75±0.64 | Medium |
|  | H<sub>2</sub>O<sub>2</sub> | 0.433 | 7.0E-06 | 2.91±0.89 | Large |
|  | Salt | 0.133 | 7.0E-06 | 3.13±0.92 | Large |
|  | F | 0.129 | 7.0E-06 | 2.23±0.79 | Large |

**Figure 1:**
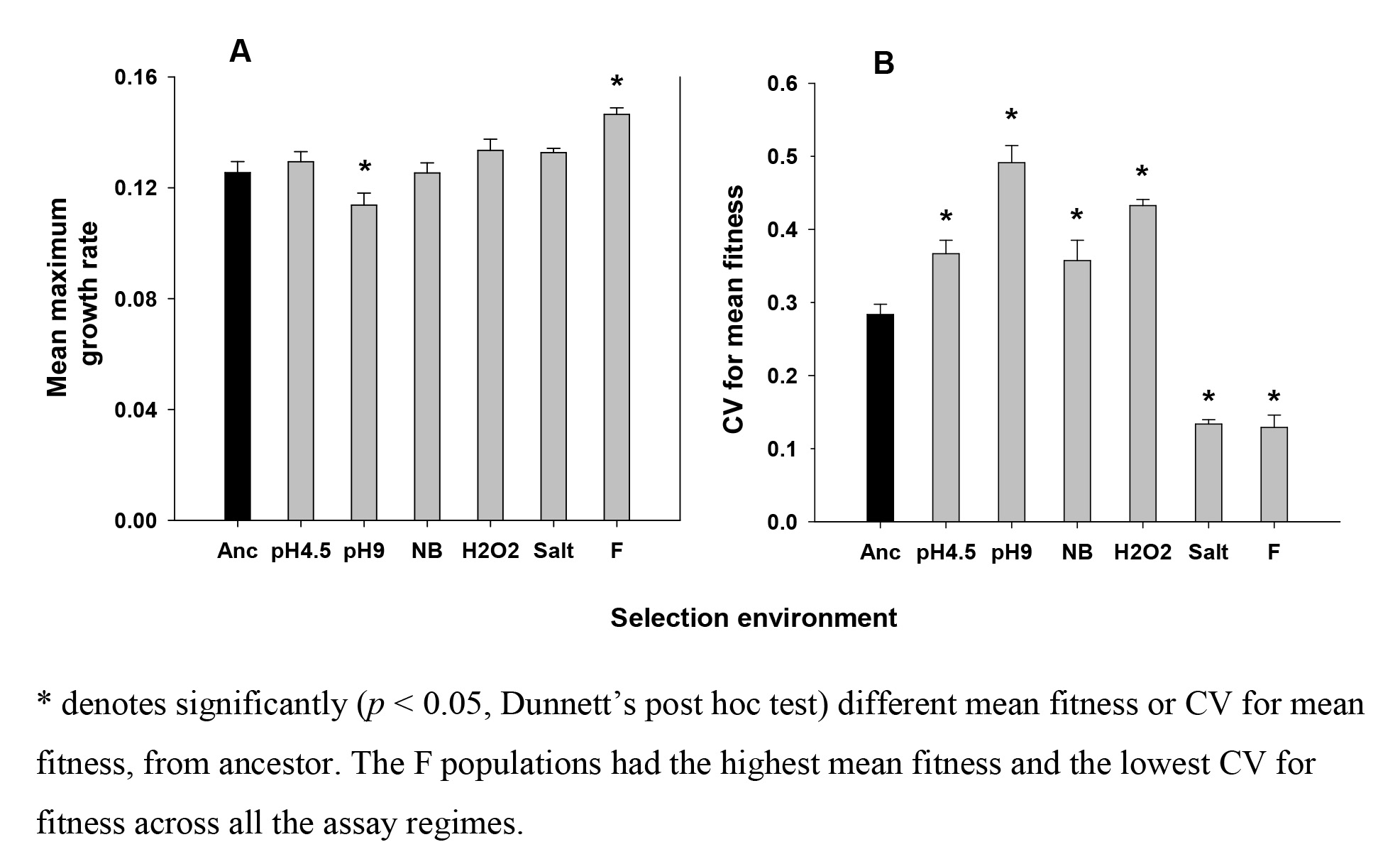
Mean fitness and CV for mean fitness for all the selection regimes across five environments. Fitness estimated as maximum slope of the growth trajectory over 24 hours. Overall mean fitness was computed for every selection regime over all assay environments. Coefficient of variation was computed for every selection regime over all the assay environments. Error bars represent SEM.

Selection also had a significant effect on the variation of fitness across all environments (*F*_6,133_ = 61.65, *p* < 0.0001). The coefficient of variation of the F populations was the lowest among all the treatments and it was significantly lower than the ancestor with a large effect size (Table 1, Fig 1B). The populations selected in salt also had significantly lower variation for fitness (compared to the ancestor) with large effect size (Table 1, Fig 1B). All the other populations had significantly greater variation in fitness than the ancestor across all environments.

To summarize, our results show that when subjected to such complex, unpredictable fluctuations for ~ 900 generations, the bacterial populations show an overall (i.e. across environments) modest increase in mean growth rate in the stresses under which they evolved (Fig 1A). To gain a more nuanced understanding of the patterns of fitness evolution, we separately examined the growth rates for each selection regime in each environment (Fig S4).

### Fluctuating environments minimize trade-offs in growth rates

Compared to the ancestors, the F populations did not show any significant loss in fitness in any of the environments (Fig 2). In contrast, populations selected in constant environments always showed a significant loss of fitness in at least one of the environments (Table S5 (A), Fig 2). All the selected populations showed significant increase in fitness in hydrogen peroxide and all of them, except F populations, show loss of fitness in salt. Interestingly, even the populations selected in salt showed reduction in fitness when assayed in salt, albeit to a lesser degree (see discussion).

**Figure 2:**
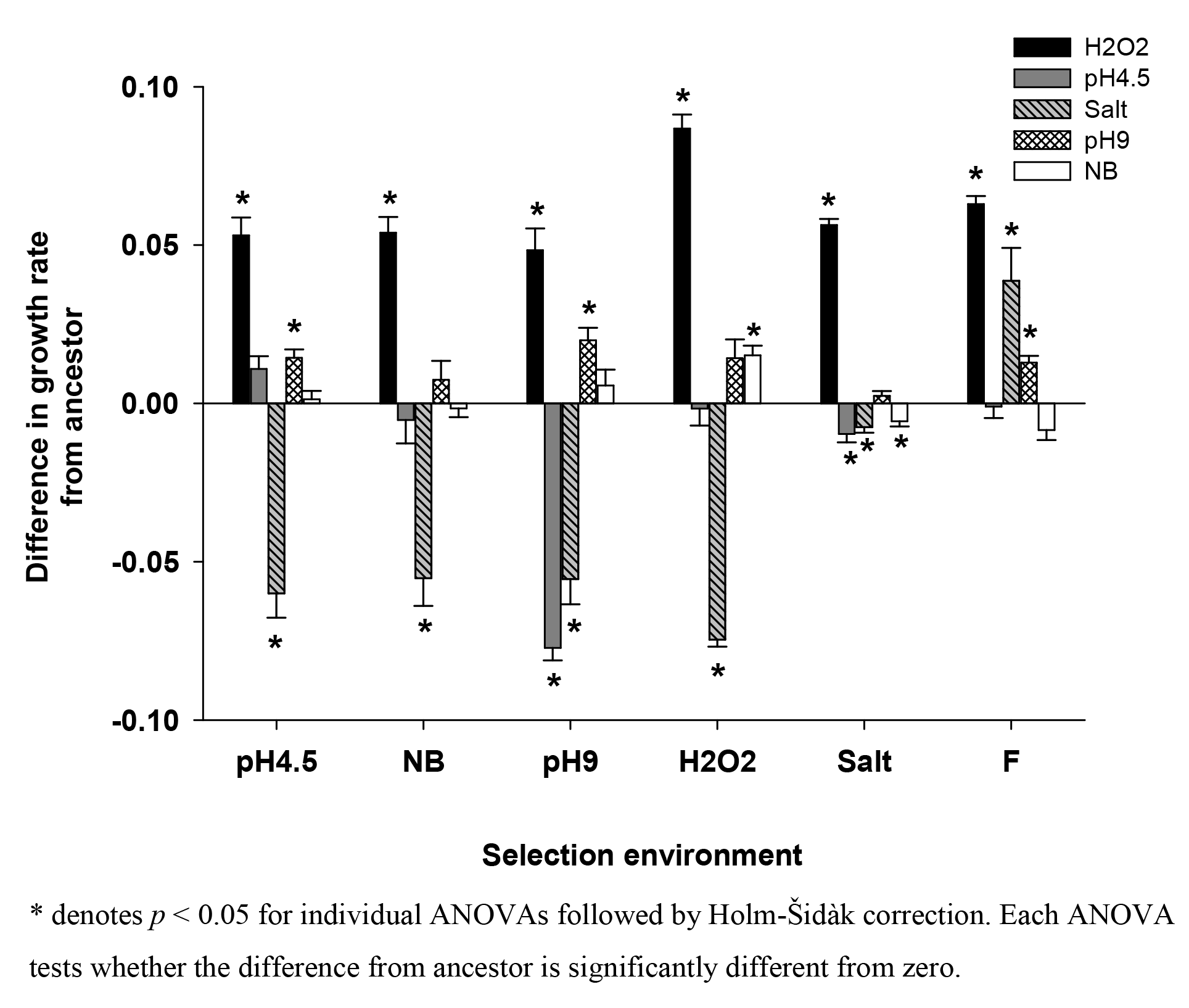
Difference (±SE) in maximum growth rate from ancestor for all the selection regimes. Difference in mean fitness (estimated as maximum growth rate) from ancestor was computed for every selection regime in every selection environment. Negative values indicate loss of fitness while positive values indicate gain of fitness, as compared to the ancestor. Every selection regime, except F, shows loss of fitness in at least one of the environments.

Taken together, these results show that fluctuating environments can minimize trade-offs across environments.

### Trade-offs can be seen across two different components of fitness

When we analyzed the mean and variance in fitness across environments, estimated as carrying capacity, the patterns in overall mean and variance were similar to those obtained with maximum growth rate (Fig 4), i.e. F populations showed the highest mean fitness with lowest variation for fitness across environments. Compared to the ancestors, F populations improved fitness in every selection environment with an exception of pH4.5 (Fig 3). Interestingly, populations selected in salt showed improved carrying capacity in salt as well as NB even though there was a significant reduction in maximum growth rate in these two assay environments. These results suggest that apart from trade-offs across environments, microbial populations can also exhibit trade-offs across different components of fitness in a given environment.

**Figure 3:**
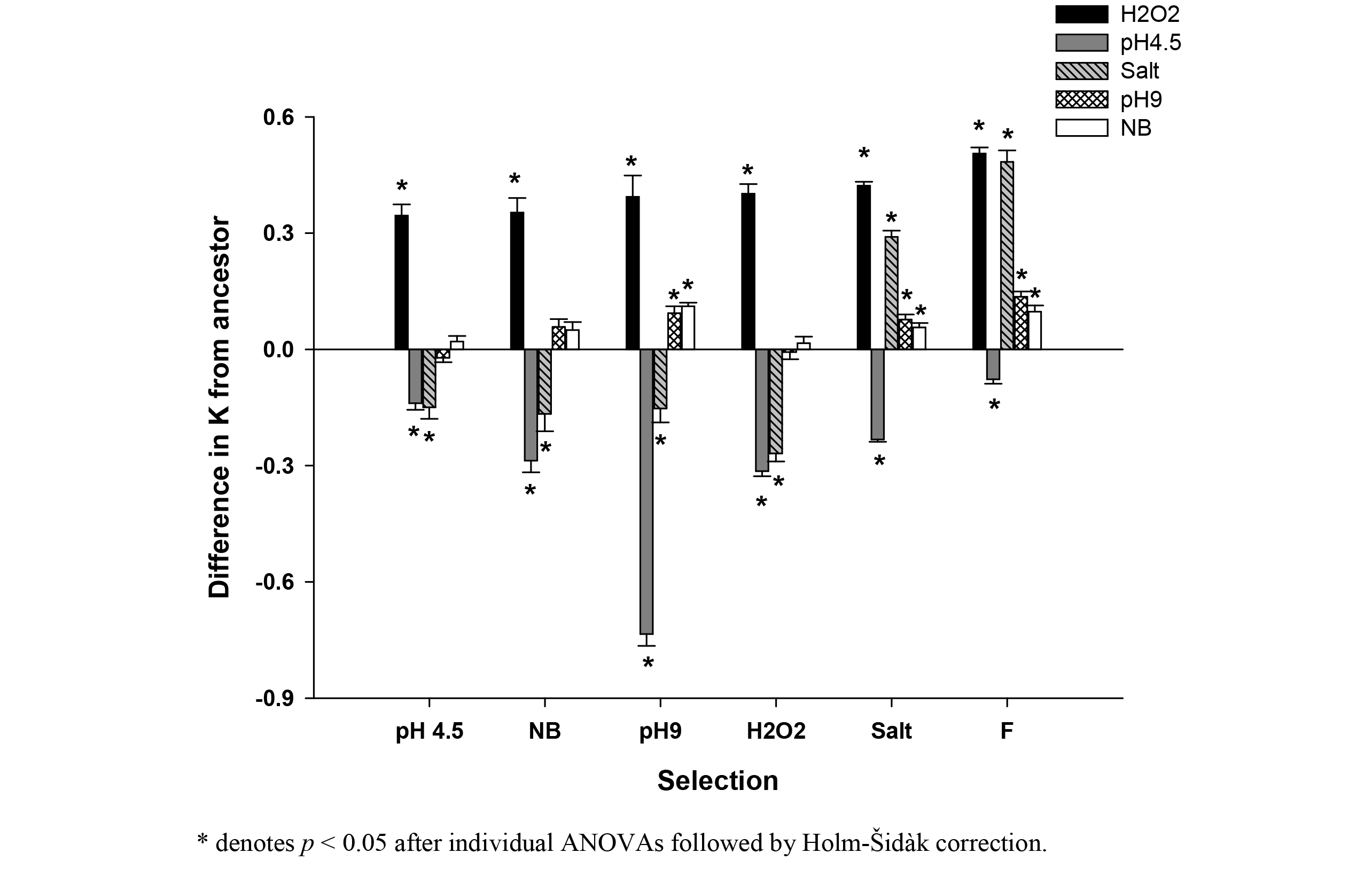
Difference (±SE) in K from ancestor for all the selection regimes. Difference in mean fitness (estimated as maximum density reached i.e. K) from ancestor was computed for every selection regime in every selection environment. Negative values indicate loss of fitness while positive values indicate gain of fitness, as compared to the ancestor. Every selection regime, shows loss of fitness in at least one of the environments.

**Figure 4:**
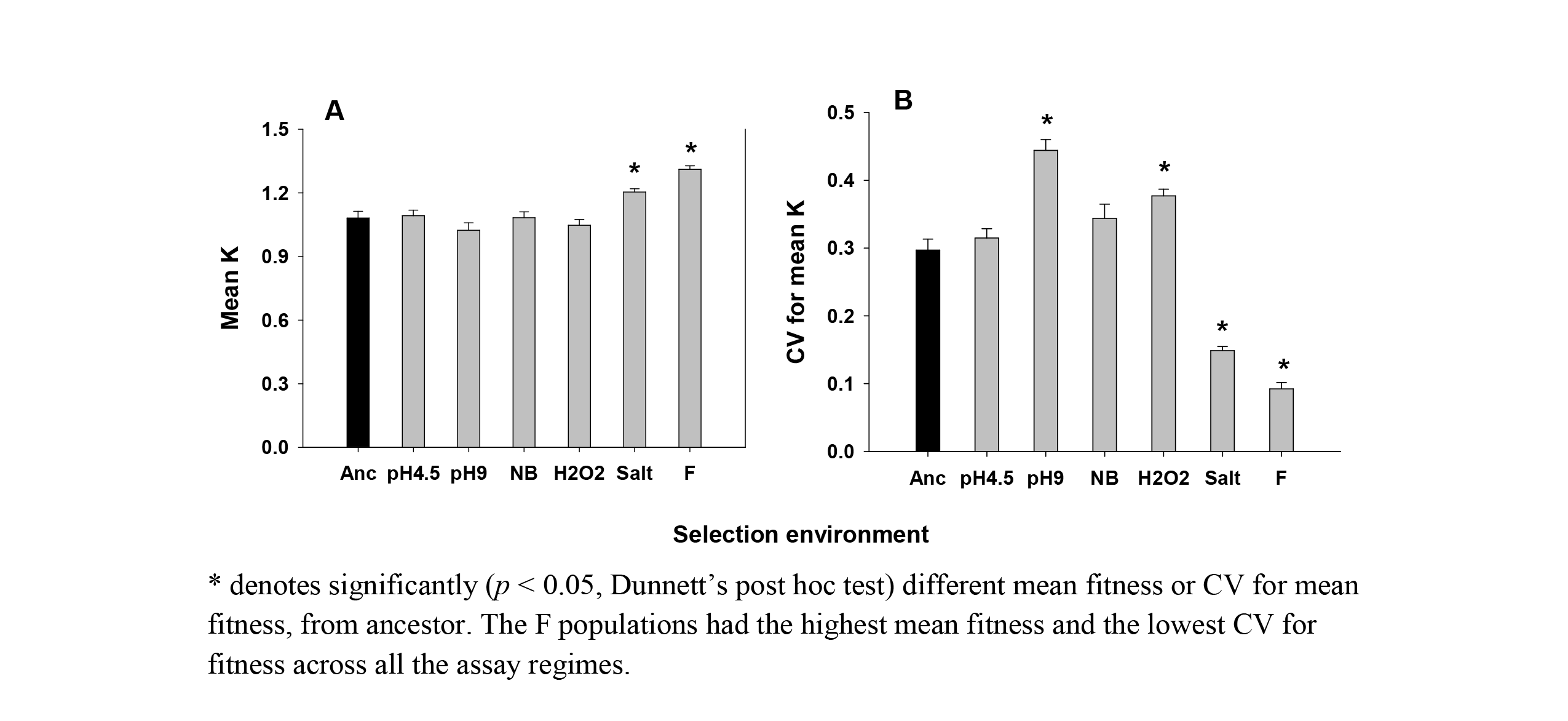
Mean *K* (±SE) and CV for mean *K* for all the selection regimes. Fitness estimated as maximum density reached (*K*) in the growth trajectory of 24 hours. Overall mean fitness was computed for every selection regime over all assay environments. Coefficient of variation (CV) was computed for every selection regime over all the assay environments. Error bars represent SEM.

## DISCUSSION

### Complex, unpredictable fluctuations select for higher overall mean fitness across environments

When subjected to predictable oscillations in a single environmental parameter over long time scales, microbial populations typically evolve to have higher fitness over the entire range of environments faced (Leroi *et al.*, 1994; Turner & Elena, 2000; Hughes *et al.*, 2007; Coffey & Vignuzzi, 2011; Alto *et al.*, 2013; Puentes-Téllez *et al.*, 2013; Condon *et al.*, 2014). However, the evolutionary outcomes become much more diverse when the environment undergoes unpredictable fluctuations and the fitness of the selected populations can show no change (Alto *et al.*, 2013), increase across the board (Turner & Elena, 2000; Cooper & Lenski, 2010; Ketola *et al.*, 2013) or an increase with respect to a few life history traits with no change (Hughes *et al.*, 2007) or decrease with respect to others (Hallsson & Björklund, 2012). To further complicate matters, natural environments typically consist of multiple parameters that can change simultaneously (Lindow & Brandl, 2003; Okafor, 2011). A previous study had reported little improvement in overall mean fitness when *E. coli* were selected in complex, unpredictable fluctuations over a short duration of ~170 generations (Karve *et al.*, 2015). However, our results show that when subjected to such complex, unpredictable fluctuations for ~ 900 generations, the bacterial populations show modest increase in fitness in the stresses under which they evolved (Fig 1A).

### Fluctuating environments minimize the variation for fitness across environments

When evolved under predictable or unpredictable temporal fluctuations, microbial populations show reduced variation for fitness over the whole range of selection environments (Kassen, 2014) Table 4.2). This is because, when the environment changes temporally, it is the geometric mean (and not the arithmetic mean) of the fitness over the entire evolutionary time that plays a more important role in determining the long-term evolutionary success (Gillespie, 1977). Since reducing the variation in a series increases its geometric mean (Orr, 2007), populations facing fluctuating environments are expected to evolve lower variation in fitness across the selection environments. This prediction is supported by empirical studies where pH or host type fluctuates across time (Kassen, 2014) Table 4.2). However, reducing the variation in fitness over different environments when multiple selection environments are changing unpredictably, is expected to be much more challenging particularly when there are negative pleiotropic interactions between the traits. Interestingly, our results show that the populations selected under complex, unpredictable fluctuations did minimize the variation in the fitness across all selection environments (Fig 1B).

The reduced fitness variation across environments in the F populations suggests the possibility of evolution of bet-hedging. However, reduced variation in growth rate and carrying capacity observed when the stresses were experienced singly, do not necessarily imply that the growth rate variations of the F populations were indeed less while they were facing unpredictably changing complex environments during the selection process. In order to establish that claim, one would be required to assay the fitness of all the populations during the process of selection itself (Collins, 2011). Although technically possible, such an exercise was not attempted here due to logistic reasons. Another way in which bet hedging is often defined in microbial evolution is as a strategy that maximizes the overall reproductive success through evolution of phenotypic heterogeneity in a formerly isogenic (and hence presumably phenotypically homogeneous) population (de Jong *et al.*, 2011). Since all the fitness assays in this study were conducted at the population level, it is difficult for us to comment on whether phenotypic heterogeneity, and hence bet-hedging across genotypes, has indeed evolved in the F populations. We note here though that one previous study looking at the fitnesses of individual cells from populations selected under complex unpredictably, fluctuating environments, suggests lack of phenotypic heterogeneity in the face of multiple novel environments (Karve *et al.*, 2016). Characterizing the individual cells of F populations across environments can be a potential way of deducing whether bet-hedging has evolved, but is out of the scope of this study.

To summarize, the present study does not allow us to conclude whether bet-hedging has evolved in the F populations were not. The significantly lower variation for fitness in F populations (Fig 1B) compared to the ancestor needs to be interpreted together with the observation that the F populations also show significantly higher overall mean fitness (Fig 1A). There are multiple ways in which these observed results can be obtained. To elucidate the matter further, we estimated the differences in fitness from the ancestors under different environments.

### Fluctuating environment minimizes trade-off

All else being equal, when an ancestral population is close to the fitness maxima in a given environment, there will be little response to selection. On the other hand when the ancestor is far away from the fitness maxima for a given environment, given the required genetic variation, improvement in fitness in the selection environment is expected, even if it is accompanied by loss of fitness in other environments. However, improvement in fitness can be constrained in case of environmental heterogeneity (temporally fluctuating or complex or both), where populations face multiple environments during selection. If an increase in fitness in one environment is accompanied by a decrease in fitness in another environment, then the presence of such trade-off structures can potentially reduce the rate of adaptation, cause no adaptation or might even lead to maladaptation for some environments. Thus, bacterial populations facing multiple environments (simultaneously or sequentially) are expected to show lesser increase in fitness as compared to that of the specialists. However, results of empirical studies, involving both predictable and unpredictable fluctuations in single environmental parameter or host, do not agree with these predictions. Populations facing multiple values of a given environmental parameter can improve fitness over all or some of the selection environments without a loss of fitness in other selection environments (Turner & Elena, 2000; Hughes *et al.*, 2007; Condon *et al.*, 2014). Our results extend this understanding to complex environments where multiple parameters fluctuate unpredictably over time. The populations selected under constant exposure to hydrogen peroxide and pH9 showed increase in fitness in the respective selection environments (Fig 2) which suggest that the ancestor was away from the fitness maxima of these two selection environments. However, this increase was accompanied by a loss of fitness in environments not experienced during selection (Fig 2). On the other hand, although the F populations showed increase in fitness for pH 9 and hydrogen peroxide, they did not lose fitness in any of the selection environments (Fig 2). In other words, the F populations not only adapted to these environments but could also by-pass the trade-offs that were experienced by the pH9 or hydrogen peroxide selected populations.

In case of pH4.5, neither F populations nor the populations selected under constant pH4.5 showed improved fitness over the ancestor (Fig 2). Thus, the ancestor was most likely well adapted to pH4.5 which is not surprising given that *Escherichia coli* are known to be well-adapted to acidic environments (reviewed in Foster, 2004). Similarly, control environment of NB^Kan^ shows no improvement in fitness, which is intuitive given the ancestral *Escherichia coli* strain K12 is expected to be well adapted to the laboratory conditions.

The populations facing complex, fluctuating environments seem to face negligible constraints while adapting to component selection environments. The extent of adaptation seems to be governed by the distance of the ancestor from fitness maxima rather than the constraints imposed by the environment. In fact in salt, F populations show gain of fitness as compared to the ancestor while populations selected in constant exposure to salt display loss of fitness (Fig 2). One possible reason for this loss of fitness in salt selected populations could be that salt induces the *rpo-S* mediated pathways in *E. coli* (Hengge-Aronis *et al.*, 1993), which in-turn can increase the rate of mutagenesis (Layton & Foster, 2003). Over generations, this can lead to an accumulation of stress-induced deleterious mutations in the genome, which might reduce the growth rate of the salt-selected populations in other environments. However, the observation that the salt populations had an appreciable growth rate increase in H_2_O_2_ (Fig 2) was not entirely consistent with this conjecture. Therefore, we investigated another way by which the poor growth rate of the salt-selected populations could be explained, namely through the evolution of a different component of fitness. For this, we studied the carrying capacity (K), estimated as the maximum density achieved by the populations during a growth curve assay (Novak *et al.*, 2006).

### Trade-off can be observed across traits for a given environment

Populations selected in salt showed increased *K* relative to the ancestor (Fig 3). This improvement in *K*, and lack of the same in maximum growth rate as a result of selection, could be due to an underlying trade-off between growth-rates and carrying capacity. Such trade-off can be brought about in more than one ways. For instance, higher concentration of salt can result in strong selection for robust membrane structures, which can be negatively correlated with the growth rate of the cells (Carlquist *et al.*, 2012). Alternatively, since our experiment involves sub-culturing every 24 hours (i.e. typically after the population has attained the stationary phase), those genotypes that are more in number at that stage can have proportionately greater representation in the next generation. This might be attained, for example, by being able to stay alive and divide at low nutrient conditions which would give a competitive advantage (Tilman *et al.*, 1981). Consistent with the observations for maximum growth rate as a proxy of fitness, even the F populations showed corresponding improvement in *K*. In the light of these results though, the choice of fitness measure demands further attention.

In many studies on microbial experimental evolution, competitive ability relative to the ancestor is used as a proxy of fitness (Leroi *et al.*, 1994; Hughes *et al.*, 2007; Alto *et al.*, 2013). This measure is preferred because it integrates over all phases of a growth cycle (i.e. lag phase, log phase etc.) and is expected to show how much better the evolved population has become in terms of evolutionarily replacing the ancestor (Kassen, 2014 Page 16). However, we refrained from using this measure in the present study. This is because change in competitive fitness compared to the ancestor is an appropriate measure in case of constant or directionally changing selection environments, where populations are adapting towards a fixed fitness peak (Collins, 2011). As opposed to this, adaptation to unpredictable environments will involve sudden changes in the underlying fitness landscape. Populations facing such environments will not show monotonic change in the relative fitness as compared to the ancestors. Thus, estimating the competitive fitness against the ancestor at any given point in the evolutionary trajectory does not give us much useful information about how much the population has really changed (Collins, 2011). Incidentally, the same argument holds for any other measurement of fitness including the ones that we have used in this study (growth rate and carrying capacity). Therefore, these two indices are perhaps better viewed as fitness-related properties of the populations and in that sense are in the same epistemological category as competitive ability vis-a-vis the ancestors. Focussing on these two fitness-properties allows us to draw some interesting conclusions. The results of maximum growth rate and *K* are comparable in case of overall mean fitness and variation for fitness (Figs 1 and 4). But this is not the case when we consider the change in fitness from ancestor (Figs 2 and 3). Populations selected in salt show reduced maximum growth rate as compared to the ancestor but significantly higher K. Similarly in pH 4.5, maximum growth rate does not evolve in comparison to ancestor in all the selected populations but *K* shows a significant reduction (Fig 2 and 3). This shows that evolutionary interpretations based on different components of fitness may or may not agree with each other and different selection environments can select for different components of fitness. This is analogous to the concept of stability properties in ecology (Grimm & Wissel, 1997) where it has been shown that evolution of one kind of stability may or may not lead to the evolution of another type (Dey *et al.*, 2008). Thus, concentrating on any one fitness property in experimental evolution studies might lead to an under-estimation of the richness of the evolutionary process.

## Conclusions

To our knowledge, this is the first study which shows that populations can improve the fitness and minimize the variation for fitness across environments when exposed to complex, unpredictable fluctuating environments. Simultaneous exposure can result in fitness increase in some selection environments without any loss of fitness in other selection environments. More importantly, our results suggest a possible explanation for the absence of trade-offs across environments in microbial populations by showing that other kind of trade-offs can exist i.e. trade-offs between different components of fitness in the same environment. Estimating different components of fitness separately might be fruitful for future experimental studies.

## Acknowledgements

We thank Dr. Manjula Reddy for providing Kanamycin resistant strain of *Escherichia coli* K12 and Somendra Singh Kharola and S. Selweshwari for their help in laboratory work. This manuscript was greatly benefitted by critical comments from Luke Holman and an anonymous reviewer. SK was supported by a Senior Research Fellowship from Council of Scientific and Industrial Research, Govt. of India. This project was supported by a grant from Department of Biotechnology, Government of India and internal funding from Indian Institute of Science Education and Research, Pune. The authors declare no conflict of interests.

## Supporting information

### S1. Details of the ancestral *Escherichia coli* population used for the selection

We used an *Escherichia coli* K12 MG1655 strain in which the lacY gene had been replaced with a Kanamycin resistance gene. Colonies of this bacterium are white coloured on MacConkey’s agar as opposed to the red coloured colonies produced by other *Escherichia coli*.

### S2. Composition of Nutrient broth

S2. Composition of Nutrient broth -
| Ingredients | Gms/litre |
| --- | --- |
| Peptic digest of animal tissue | 5.00 |
| Sodium chloride | 5.00 |
| Beef extract | 1.50 |
| Yeast extract | 1.50 |

Final pH (at 25<sup>0</sup>C) 7.4 ± 0.2

For making NB^Kan^, 0.05 mg/ml of Kanamycin was added to the above mixture after autoclaving and cooling.

We added 2g/100ml of agar to the above mixture before autoclaving, to make Nutrient Agar^Kan^.

### S3. Details of all the selection regimes

Component space for fluctuating selection regime –

a. Salt – 0.5 g%, 2 g%, 3 g%, 3.5 g%, 4 g%, 4.5 g%, 5 g%
b. pH – 4.5, 5, 7, 8.5, 9
c. Hydrogen peroxide – 2.4 mM, 2.9 mM, 3.4 mM, 3.9 mM

Selection regime faced by F populations for 100 days

| Transfer # | Combination # | Salt g% | pH | H<sub>2</sub>O<sub>2</sub> mM |
| --- | --- | --- | --- | --- |
| 1 | 39 | 5 | 7 | 2.9 |
| 2 | 59 | 5 | 7 | 3.9 |
| 3 | 2 | 3.5 | 5 | 0 |
| 4 | 46 | 3.5 | 7 | 3.4 |
| 5 | 23 | 5 | 10 | 0 |
| 6 | 10 | 4.5 | 4.5 | 0 |
| 7 | 62 | 0.5 | 8.5 | 3.9 |
| 8 | 1 | 3 | 5 | 0 |
| 9 | 19 | 3 | 10 | 0 |
| 10 | 9 | 4 | 4.5 | 0 |
| 11 | 9 | 4 | 4.5 | 0 |
| 12 | 30 | 0.5 | 5 | 2.4 |
| 13 | 7 | 3 | 4.5 | 0 |
| 14 | 39 | 5 | 7 | 2.9 |
| 15 | 45 | 3 | 7 | 3.4 |
| 16 | 32 | 0.5 | 8.5 | 2.4 |
| 17 | 44 | 2 | 7 | 3.4 |
| 18 | 58 | 4.5 | 7 | 3.9 |
| 19 | 60 | 0.5 | 5 | 3.9 |
| 20 | 36 | 3.5 | 7 | 2.9 |
| 21 | 9 | 4 | 4.5 | 0 |
| 22 | 45 | 3 | 7 | 3.4 |
| 23 | 25 | 3 | 7 | 2.4 |
| 24 | 12 | 2 | 8.5 | 0 |
| 25 | 8 | 3.5 | 4.5 | 0 |
| 26 | 11 | 5 | 4.5 | 0 |
| 27 | 25 | 3 | 7 | 2.4 |
| 28 | 34 | 2 | 7 | 2.9 |
| 29 | 26 | 3.5 | 7 | 2.4 |
| 30 | 26 | 3.5 | 7 | 2.4 |
| 31 | 53 | 0.5 | 9 | 3.4 |
| 32 | 18 | 2 | 9 | 0 |
| 33 | 55 | 3 | 7 | 3.9 |
| 34 | 43 | 0.5 | 9 | 2.9 |
| 35 | 27 | 4 | 7 | 2.4 |
| 36 | 24 | 2 | 7 | 2.4 |
| 37 | 48 | 4.5 | 7 | 3.4 |
| 38 | 1 | 3 | 5 | 0 |
| 39 | 21 | 4 | 9 | 0 |
| 40 | 14 | 3.5 | 8.5 | 0 |
| 41 | 48 | 4.5 | 7 | 3.4 |
| 42 | 63 | 0.5 | 9 | 3.9 |
| 43 | 30 | 0.5 | 5 | 2.4 |
| 44 | 49 | 5 | 7 | 3.4 |
| 45 | 30 | 0.5 | 5 | 2.4 |
| 46 | 21 | 4 | 9 | 0 |
| 47 | 46 | 3.5 | 7 | 3.4 |
| 48 | 62 | 0.5 | 8.5 | 3.9 |
| 49 | 17 | 5 | 8.5 | 0 |
| 50 | 49 | 5 | 7 | 3.4 |
| 51 | 41 | 0.5 | 4.5 | 2.9 |
| 52 | 63 | 0.5 | 9 | 3.9 |
| 53 | 25 | 3 | 7 | 2.4 |
| 54 | 25 | 3 | 7 | 2.4 |
| 55 | 28 | 4.5 | 7 | 2.4 |
| 56 | 16 | 4.5 | 8.5 | 0 |
| 57 | 46 | 3.5 | 7 | 3.4 |
| 58 | 55 | 3 | 7 | 3.9 |
| 59 | 31 | 0.5 | 4.5 | 2.4 |
| 60 | 3 | 4 | 5 | 0 |
| 61 | 30 | 0.5 | 5 | 2.4 |
| 62 | 5 | 5 | 5 | 0 |
| 63 | 42 | 0.5 | 8.5 | 2.9 |
| 64 | 28 | 4.5 | 7 | 2.4 |
| 65 | 2 | 3.5 | 5 | 0 |
| 66 | 47 | 4 | 7 | 3.4 |
| 67 | 1 | 3 | 5 | 0 |
| 68 | 24 | 2 | 7 | 2.4 |
| 69 | 22 | 4.5 | 9 | 0 |
| 70 | 38 | 4.5 | 7 | 2.9 |
| 71 | 7 | 3 | 4.5 | 0 |
| 72 | 18 | 2 | 9 | 0 |
| 73 | 46 | 3.5 | 7 | 3.4 |
| 74 | 2 | 3.5 | 5 | 0 |
| 75 | 0 | 2 | 5 | 0 |
| 76 | 11 | 5 | 4.5 | 0 |
| 77 | 40 | 0.5 | 5 | 2.9 |
| 78 | 24 | 2 | 7 | 2.4 |
| 79 | 25 | 3 | 7 | 2.4 |
| 80 | 37 | 4 | 7 | 2.9 |
| 81 | 56 | 3.5 | 7 | 3.9 |
| 82 | 21 | 4 | 9 | 0 |
| 83 | 43 | 0.5 | 9 | 2.9 |
| 84 | 62 | 0.5 | 8.5 | 3.9 |
| 85 | 53 | 0.5 | 9 | 3.4 |
| 86 | 4 | 4.5 | 5 | 0 |
| 87 | 63 | 0.5 | 9 | 3.9 |
| 88 | 42 | 0.5 | 8.5 | 2.9 |
| 89 | 62 | 0.5 | 8.5 | 3.9 |
| 90 | 33 | 0.5 | 9 | 2.4 |
| 91 | 48 | 4.5 | 7 | 3.4 |
| 92 | 43 | 0.5 | 9 | 2.9 |
| 93 | 50 | 0.5 | 5 | 3.4 |
| 94 | 11 | 5 | 4.5 | 0 |
| 95 | 8 | 3.5 | 4.5 | 0 |
| 96 | 35 | 3 | 7 | 2.9 |
| 97 | 17 | 5 | 8.5 | 0 |
| 98 | 59 | 5 | 7 | 3.9 |
| 99 | 11 | 5 | 4.5 | 0 |
| 100 | 15 | 4 | 8.5 | 0 |

Transfer #5 and #9 comprised of pH 10 which was initially part of the fluctuating component space. Both the environments lead to extinction and hence the component space was modified to the present state.

Assay concentrations for the five component environments were as follows –

1. Salt – 5g% of sodium chloride
2. Acidic pH – pH 4.5
3. Basic pH – pH 9
4. Hydrogen peroxide – 3.9 mM

### S4. Growth rates for each selected population in every assay environment

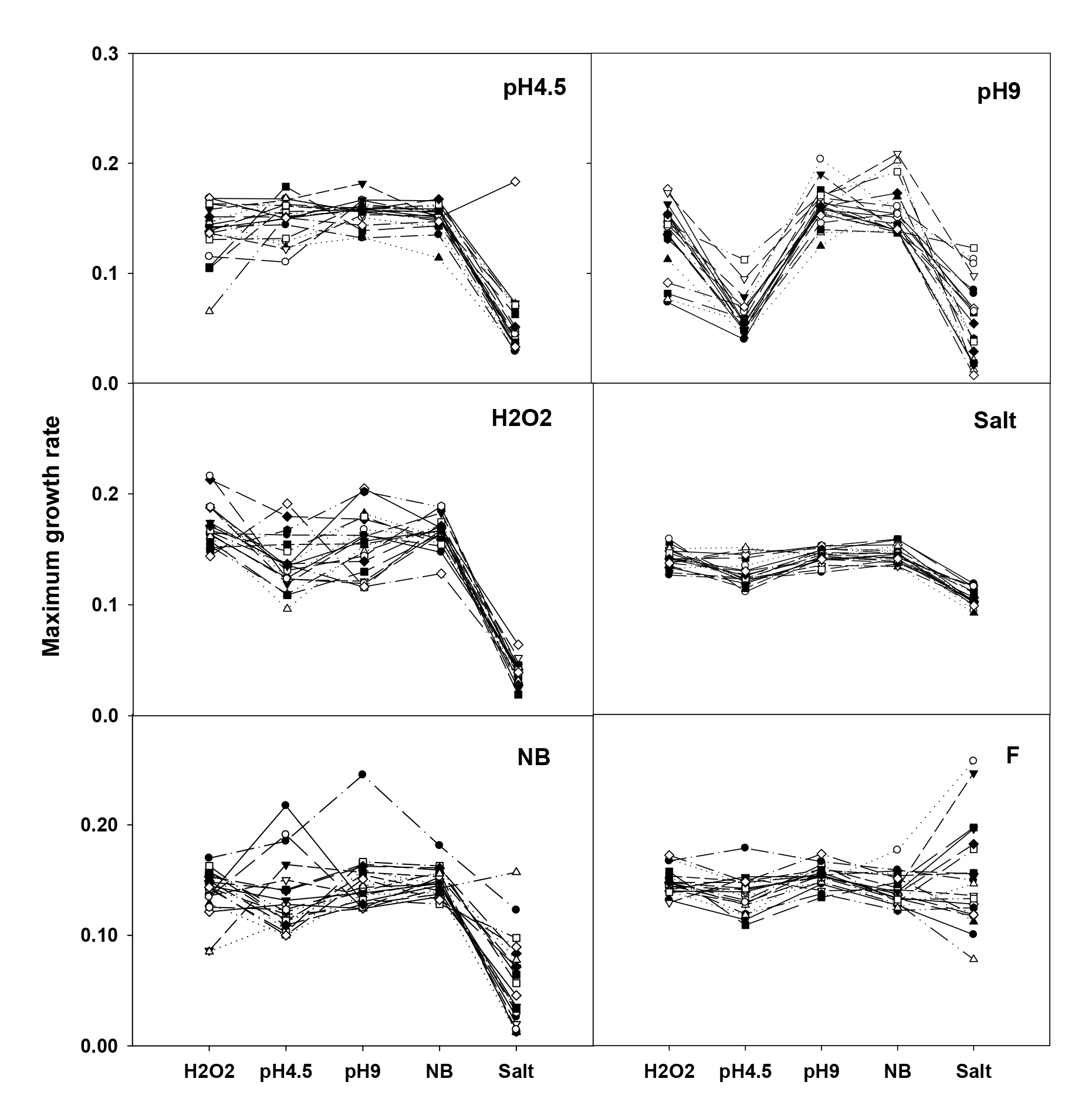

Fitness estimated as maximum slope of the growth trajectory over 24 hours. Each panel shows maximum growth rate of 20 replicate populations of a given selection regime, across all the assay environments. Every point is an average of two measurements for a given combination of selection and assay environment.

### Table S5: Difference from ancestor was computed for all the selection regimes in all the assay environments

Holm – Šidàk corrected *p* values for 30 t-tests are given for (A) maximum growth rate and (B) maximum density achieved. *p* < 0.05 denotes significant difference from ancestor which is accompanied by a sign in the bracket denoting direction of change. ‘-’ represents decrease while ‘+’ represents increase in fitness from ancestor.

| Assay environment ↓ | Selection environment → |  |  |  |  |  |
| --- | --- | --- | --- | --- | --- | --- |
|  | pH4.5 | pH9 | NB | H<sub>2</sub>O<sub>2</sub> | Salt | F |
| **pH4.5** | 0.14 | **1.3E-12(-)** | 0.96 | 0.95 | **0.02(-)** | 0.78 |
| **pH9** | **0.0004(+)** | **0.001(+)** | 0.84 | 0.22 | 0.71 | **0.0001(+)** |
| **NB** | 0.95 | 0.85 | 0.97 | **0.001(+)** | **0.02(-)** | 0.14 |
| **H<sub>2</sub>O<sub>2</sub>** | **3.1E-07(+)** | **2.4E-05(+)** | **3.1E-08(+)** | **1.1E-12(+)** | **6.4E-15(+)** | **2.2E-14(+)** |
| **Salt** | **5.4E-06 (-)** | **2.5E-05(-)** | **9.6E-05(-)** | **6.7E-15(-)** | **0.006(-)** | **0.01(+)** |

(B)
| Assay environment ↓ | Selection environment → |  |  |  |  |  |
| --- | --- | --- | --- | --- | --- | --- |
|  | pH4.5 | pH9 | NB | H<sub>2</sub>O<sub>2</sub> | Salt | F |
| **pH4.5** | **1.9E-06(-)** | **2.6E-14(-)** | **2E-07(-)** | **2.7E-14(-)** | **6.4E-15(-)** | **3E-05(-)** |
| **pH9** | 0.25 | **0.0004(+)** | 0.06 | 0.71 | **0.0001(+)** | **8.8E-08(+)** |
| **NB** | 0.44 | **5.3E-09(+)** | 0.12 | 0.6 | **0.0011(+)** | **0.0001(+)** |
| **H<sub>2</sub>O<sub>2</sub>** | **5.3E-09(+)** | **1.3E-05(+)** | **2.8E-07(+)** | **3.7E-11(+)** | **6.7E-15(+)** | **6.2E-15(+)** |
| **Salt** | **0.0007(-)** | **0.003(-)** | **0.01(-)** | **1.6E-09(-)** | **6.9E-12(+)** | **3.3E-11(+)** |

## References

Abdi, H. 2010. Holm’s sequential Bonferroni procedure. Encyclopedia of Research Design. N. Salkind, Sage Thousand Oaks. California: 1–8.

Acar, M., Mettetal, J.T.& van Oudenaarden, A. 2008. Stochastic switching as a survival strategy in fluctuating environments. Nat. Genet. 40: 471–475.

Alto, B.W., Wasik, B.R., Morales, N.M. & Turner, P.E. 2013. Stochastic temperatures impede RNA virus adaptation. Evolution 67: 969–979.

Ancel, L.W. 1999. A quantitative model of the Simpson-Baldwin effect. J. Theor. Biol. 196: 197–209.

Anderson, J.T., Lee, C.-R. & Mitchell-Olds, T. 2014. Strong selection genome-wide enhances fitness trade-offs across environments and episodes of selection. Evolution 68: 16–31.

Barrett, R.D.H., MacLean, R.C. & Bell, G. 2005. Experimental evolution of Pseudomonas fluorescens in simple and complex environments. Am. Nat. 166: 470–480.

Beaumont, H.J.E., Gallie, J., Kost, C., Ferguson, G.C. & Rainey, P.B. 2009. Experimental evolution of bet hedging. Nature 462: 90–93.

Bennett, Albert F., Lenski, R.E. & Mittler, J.E. 1992. Evolutionary adaptation to temperature. I. Fitness responses of Escherichia coli to changes in its thermal environment. Evolution 46: 16–30.

Bennett, A.F. & Lenski, R.E. 1997. Phenotypic and evolutionary adaptation of a model bacterial system to stressful thermal environments. In: Environmental Stress, Adaptation and Evolution (R. Bijlsma & V. Loeschcke, eds). Birkhäuser Verlag, Basel.

Carlquist, M., Fernandes, R.L., Helmark, S., Heins, A.-L., Lundin, L., Sørensen, S.J., Gernaey, K.V. & Lantz, A.E. 2012. Physiological heterogeneities in microbial populations and implications for physical stress tolerance. Microb. Cell. Fact. 11: 94.

Chevin, L.-M., Lande, R. & Mace, G.M. 2010. Adaptation, plasticity, and extinction in a changing environment: towards a predictive theory. PLoS Biol. 8: e1000357.

Coffey, L.L. & Vignuzzi, M. 2011. Host alternation of chikungunya virus increases fitness while restricting population diversity and adaptability to novel selective pressures. J. Virol. 85: 1025–1035.

Cohan, F.M. 2005. Periodic selection and ecological diversity in bacteria. In: Selective Sweep (D. Nurminsky, ed., pp. 78–93. Eurekah.com and Kluwer Academic / Plenum Publishers., Georgetown/ New York.

Cohen, J. 1988. Statistical Power Analysis for the Behavioral Sciences. Lawrence Erlbaum Associates, USA.

Collins, S. 2011. Many possible worlds: expanding the ecological scenarios in experimental evolution. Evol. Biol. 38: 3–14.

Condon, C., Cooper, B.S., Yeaman, S. & Angilletta, M.J. 2014. Temporal variation favors the evolution of generalists in experimental populations of Drosophila melanogaster. Evolution 68: 720–728.

Cooper, T.F. & Lenski, R.E. 2010. Experimental evolution with E. coli in diverse resource environments. I. Fluctuating environments promote divergence of replicate populations. BMC Evol. Biol. 10: 1–11.

de Jong, I.G., Haccou, P. & Kuipers, O.P. 2011. Bet hedging or not? A guide to proper classification of microbial survival strategies. Bioessays 33: 215–223.

Devilly, G.J. 2004. The effect size generator for Windows, Version 2.3 (computer programme), Swinburne University. Australia.

Dey, S., Prasad, N.G., Shakarad, M. & Joshi, A. 2008. Laboratory evolution of population stability in Drosophila: constancy and persistence do not necessarily coevolve. J. Anim. Ecol. 77: 670–677.

Foster, J.W. 2004. Escherichia coli acid resistance: tales of an amateur acidophile. Nat. Rev. Microbiol. 2: 898–907.

Gillespie, J.H. 1977. Natural selection for variances in offspring numbers: a new evolutionary principle. Am. Nat. 111: 1010–1014.

Graham, J.K., Smith, M.L. & Simons, A.M. 2014. Experimental evolution of bet hedging under manipulated environmental uncertainty in Neurospora crassa. Proc. R. Soc. Lond., Ser. B: Biol. Sci. 281: 20140706.

Grimm, V. & Wissel, C. 1997. Babel, or the ecological stability discussions: an inventory and analysis of terminology and a guide for avoiding confusion. Oecologia 109: 323–334.

Hallsson, L.R. & Björklund, M. 2012. Selection in a fluctuating environment leads to decreased genetic variation and facilitates the evolution of phenotypic plasticity. J. Evol. Biol. 25: 1275–1290.

Hengge-Aronis, R., Lange, R., Henneberg, N. & Fischer, D. 1993. Osmotic regulation of rpoS-dependent genes in Escherichia coli J. Bacteriol. 175: 259–265.

Holman, L. 2015. Bet hedging via multiple mating: A meta-analysis. Evolution 70: 62–71.

Hughes, B.S., Cullum, A.J. & Bennett, A.F. 2007. An experimental evolutionary study on adaptation to temporally fluctuating pH in Escherichia coli. Physiol. Biochem. Zool. 80: 406–421.

Jasmin, J.-N. & Kassen, R. 2007. Evolution of a single niche specialist in variable environments. Proc. R. Soc. B 274: 2761–2767.

Karve, S.M., Daniel, S., Chavhan, Y.D., Anand, A., Kharola, S.S. & Dey, S. 2015. Escherichia coli populations in unpredictably fluctuating environments evolve to face novel stresses through enhanced efflux activity. J. Evol. Biol. 28: 1131–1143.

Karve, S.M., Tiwary, K., Selveshwari, S. & Dey, S. 2016. Environmental fluctuations do not select for increased variation or population-based resistance in Escherichia coli. J. Biosci. 41: 39–49.

Kassen, R. 2014. Experimental evolution and the nature of biodiversity. Roberts and Company Inc., Greenwood Village, CO.

Ketola, T., Mikonranta, L., Zhang, J., Saarinen, K., Örmälä, A.-M., Friman, V.-P., Mappes, J. & Laakso, J. 2013. Fluctuating temperature leads to evolution of thermal generalism and preadaptation to novel environments Evolution 67: 2936–2944.

Layton, J.C. & Foster, P.L. 2003. Error-prone DNA polymerase IV is controlled by the stress-response sigma factor, RpoS, in Escherichia coli Mol. Microbiol. 50: 549–561.

Lenski, R.E. 2004. Phenotypic and genotypic evolution during a 20,000-generation experiment with the bacterium Escherichia coli. Plant Breed. Rev. 24: 225–265.

Leroi, A.M., Lenski, R.E. & Bennett, A.F. 1994. Evolutionary adaptation to temperature. III. Adaptation of Escherichia coli to a temporally varying environment. Evolution 48: 1222–1229.

Levins, R. 1968. Evolution in changing environments: some theoretical explorations. Princeton University Press, Princeton.

Lindow, S.E. & Brandl, M.T. 2003. Microbiology of the phyllosphere. Appl. Environ. Microbiol. 69: 1875–1883.

Novak, M., Pfeiffer, T., Lenski, R.E., Sauer, U. & Bonhoeffer, S. 2006. Experimental tests for an evolutionary trade-off between growth rate and yield in E. coli. Am. Nat. 168: 242–251.

Okafor, N. 2011. Environmental Microbiology of Aquatic and Waste Systems. Springer Science & Business Media, Berlin.

Orr, H.A. 2007. Absolute fitness, relative fitness, and utility. Evolution 61: 2997–3000.

Puentes-Téllez, P.E., Hansen, M.A., Sørensen, S.J. & van Elsas, J.D. 2013. Adaptation and heterogeneity of Escherichia coli MC1000 growing in complex environments. Appl. Environ. Microbiol. 79: 1008–1017.

Ratcliff, W.C. & Denison, R.F. 2010. Individual-level bet hedging in the bacterium Sinorhizobium meliloti Curr. Biol. 20: 1740–1744.

Reboud, X. & Bell, G. 1997. Experimental evolution in Chlamydomonas. III. Evolution of specialist and generalist types in environments that vary in space and time. Heredity 78: 507–514.

Roff, D.A. & Fairbairn, D.J. 2007. The evolution of trade-offs: where are we? J. Evol. Biol. 20: 433–447.

Scheiner, S.M. 2002. Selection experiments and the study of phenotypic plasticity. J. Evol. Biol. 15: 889–898.

Stearns, S.C. 1989. Trade-offs in life-history evolution. Funct. Ecol. 3: 259–268.

Tilman, D., Mattson, M. & Langer, S. 1981. Competition and nutrient kinetics along a temperature gradient: an experimental test of a mechanistic approach to niche theory. Limnol. Oceanogr. 26: 1020–1033.

Turner, P.E. & Elena, S.F. 2000. Cost of host radiation in an RNA virus. Genetics 156: 1465–1470.

Veening, J.W., Stewart, E.J., Berngruber, T.W., Taddei, F., Kuipers, O.P. & Hamoen, L.W.2008. Bet-hedging and epigenetic inheritance in bacterial cell development. Proc. Natl. Acad. Sci. USA 105: 4393–4398.

Whitlock, M.C. 1996. The Red Queen beats the Jack-Of-All-Trades: the limitations on the evolution of phenotypic plasticity and niche breadth. Am. Nat. 148: S65–S77.

Zar, J.H. 1999. Biostatistical Analysis. Prentice Hall, New Jersey.

